# Impaired cortical encoding of prosodic prominence in single multisyllabic words for adults with dyslexia

**DOI:** 10.64898/2026.09.25.754341

**Authors:** Mahmoud Keshavarzi, Brian C. J. Moore, Usha Goswami

## Abstract

Developmental dyslexia is associated with impaired cortical tracking of continuous speech and difficulties in perceiving within-word prosodic structure in children and adults. For adults and continuous speech, both low-frequency delta- and theta-band impairments and high-frequency gamma- and beta-band differences are found. To date, perceptual difficulties with intra-word prosody have not been studied in direct relation to neural speech encoding. Here neurotypical adults and adults with dyslexia listened to multisyllabic words. EEG was recorded, and cortical encoding was quantified using temporal response function (TRF) modelling. Four speech representations were explored: lexical stress pattern (intra-word prosodic prominence), the low-frequency speech envelope, word onsets and syllable onsets. Adults with dyslexia showed significantly reduced encoding of intra-word prosodic prominence (delta band) and of the speech envelope (delta and theta bands). In contrast, encoding of word and syllable onsets was preserved. Pinpointing delta-band cortical tracking as central to encoding intra-word prosodic prominence opens new avenues for research.

## Introduction

Successful speech perception depends on the accurate cortical transformation of continuously varying acoustic signals into meaningful linguistic representations. Speech contains temporal structure across multiple timescales, from relatively slow prosodic fluctuations to faster syllabic and phonemic fluctuations, and cortical activity dynamically tracks these temporal regularities during listening. Electrophysiological studies using electroencephalography (EEG), magnetoencephalography (MEG), and electrocorticography (ECoG) have demonstrated robust alignment between low-frequency cortical activity and the temporal structure of speech (Lalor & Foxe, 2010; Ding & Simon, 2012; Peelle et al., 2013; Ding & Simon, 2014). This neural tracking is particularly prominent in the delta (∼0.5–4 Hz) and theta (∼4–8 Hz) frequency ranges, which are associated with timescales relevant to prosodic and syllabic structure, respectively (Ghitza & Greenberg, 2009; Giraud & Poeppel, 2012; Gross et al., 2013). For adults, low-frequency cortical speech tracking is influenced not only by acoustic structure but also by speech intelligibility and selective attention, indicating that it reflects both sensory and higher-level aspects of speech processing (Peelle et al., 2013; Doelling et al., 2014). Low-frequency cortical tracking is present from infancy, where it shows clear developmental relationships with subsequent language acquisition (Attaheri et al., 2022, 2024). During childhood language development, low-frequency cortical tracking continues to play a critical role in individual differences. For example, delta band cortical tracking is selectively impaired for children with phonological impairments (such as children with developmental dyslexia, Power et al., 2013, 2016, Keshavarzi et al., 2022a; Molinaro et al., 2016; and children with developmental language disorder (DLD, Keshavarzi et al., 2026a).

The potential importance of slow temporal information for language development is evident before birth. The foetus can hear low-pass-filtered speech in the womb (Gerhardt & Abrams, 1996). Low-filtered speech retains sufficient prosodic characteristics that newborn infants can distinguish their native language (e.g., French) from a non-native language (e.g., Russian, Mehler et al., 1988), suggestive of low-frequency tracking in utero. A connection between the fidelity of delta-band cortical tracking during infancy and the encoding of continuous speech has indeed been demonstrated (Attaheri et al., 2022). However, to date, only speech modelling studies based on the amplitude envelope provide evidence that this association involves prosody (Leong et al., 2014; Leong & Goswami, 2015; Leong et al., 2017). These envelope-based modelling studies show that a unique acoustic statistic, the phase relationship between amplitude modulations at a delta-band rate (∼2 Hz) and amplitude modulations at a theta-band rate (∼5 Hz), govern the perception of syllable stress patterns in spoken English. When modulation peaks in both bands are temporally aligned, a strong syllable is perceived. When a modulation peak in the theta band is aligned with a modulation trough in the delta band, a weak syllable is perceived. The possible links between delta- and theta-band cortical encoding and the accuracy of prosodic perception have not yet been studied and are explored here for adults with and without dyslexia.

Given the central role of temporal speech encoding in constructing phonological representations, disruption of neural tracking in different frequency bands has been proposed to contribute to developmental dyslexia (Lallier et al., 2017). Developmental dyslexia is characterised by persistent difficulties in word reading despite adequate intelligence and educational opportunity (Lyon et al., 2003). Although its behavioural manifestations are most apparent during reading and writing, converging evidence indicates that dyslexia is also associated with atypical auditory and spoken-language processing (Shaywitz & Shaywitz, 2003; Goswami, 2022a). According to Temporal Sampling (TS) theory (Goswami, 2011), the phonological difficulties found in developmental dyslexia arise from atypical neural sampling of slower amplitude modulations in speech. Impaired alignment of delta- and theta-band cortical activity with prosodic and syllabic structure has been proposed to degrade the development of phonological representations from infancy onwards (Goswami, 2022b). These early-altered representations subsequently affect the development of all levels of linguistic phonology during language acquisition. For example, the difficulty with phonemic structure that is observed for older children and adults with dyslexia is thought to result from these early-altered representations, as phonemes are identified more easily in stressed syllables (Greenberg & Arai, 2004).

Consistent with TS theory, EEG and MEG studies using both children and adults with dyslexia have shown reduced or altered neural alignment to the temporal structure of speech at the delta- and theta-band timescales important for representing prosodic and syllabic structure, both during continuous natural speech listening and for rhythmic speech paradigms using single-syllable stimuli (Power et al., 2013, 2016; Molinaro et al., 2016; Destoky et al., 2020, 2022; Lizarazu et al., 2021; Mandke et al., 2022; Keshavarzi et al., 2022a,b, 2026b,c). These encoding differences have been demonstrated using complementary measures of neural speech tracking, including cortical phase entrainment, speech–brain coherence, and temporal response function (TRF) modelling, suggesting that they are not specific to a single measure.

Behaviourally, both children (Goswami et al., 2010) and adults (Leong et al., 2011) with developmental dyslexia exhibit difficulties regarding prosodic perception at the single word level. Behavioural studies have shown that adults with dyslexia are less accurate than age-matched controls at deciding whether two items like the correctly stressed “DIFF-ic-ult-y” and mis-stressed “di-FFIC-ul-ty” sound the same. They are also less accurate when judging whether different multisyllabic words, like “difficulty” and “maternity”, share the same stress pattern (Leong et al., 2011). English is a free-stressed language, and prosodic prominence may occur on different syllables, falling at different positions in different words. For example, the typical stress pattern is “orNATE” for the isolated word, but shifts in a phrase like “ORnate BALcony”, making it necessary to track prosodic prominence dynamically over time. It remains unknown how prosodic prominence within individual words is represented at the cortical level. The current study focused on isolated three- and four-syllable English words in which the primary stress occurred on the first, second, or third syllable, seeking to examine the cortical encoding of intra-word prosodic prominence for adults with and without dyslexia. Our expectation was that prosodic prominence within individual words would be represented atypically in the cortex of adults with dyslexia.

Most previous neurophysiological studies of cortical speech encoding for adults with dyslexia have used continuous speech (Lehongre et al., 2013; Lizarazu et al., 2021; Keshavarzi et al., 2026b) or speech-in-noise (Vander Ghinst et al., 2021) or extended rhythmic stimuli (Keshavarzi et al., 2026c). Cortical responses during continuous speech encoding reflect not only bottom-up encoding of the acoustic and prosodic structure of the speech signal, but also interactions with higher-level linguistic and cognitive processes. These include lexical-semantic processing, syntactic integration, contextual prediction, and working memory. Consequently, it remains unclear whether reduced low-frequency cortical speech tracking for dyslexic adults reflects an impairment in the encoding of the speech signal itself or is influenced by the additional contextual and integrative demands of continuous speech. Isolated spoken words provide a useful means of addressing this question as they preserve the acoustic, syllabic, and lexical-prosodic structure of speech while substantially reducing sentence-level contextual information and the associated demands for integration across extended timescales. Demonstrating atypical cortical encoding under these conditions would therefore indicate that the neural differences observed for those with dyslexia occur even in the absence of sentence-level contextual information.

Beyond the low-frequency speech envelope, isolated words also contain temporally precise linguistic features that contribute to spoken-word recognition. Word onsets mark the beginning of lexical units, syllable onsets provide temporal landmarks within words, and lexical stress patterns specify the relative prosodic prominence of individual syllables. Lexical stress is of particular interest here because of TS theory, and because it provides prosodic information that contributes to lexical segmentation and spoken-word recognition (Cutler & Norris, 1988; Cooper et al., 2002). Linguists conventionally code a syllable that receives primary stress in a multisyllabic word as 2, a syllable that receives no stress as 0, and a syllable that receives secondary stress as 1. Using this system, a word like ‘caterpillar’ would be coded as 2010. In the current study, syllables receiving primary stress were coded 1 and syllables receiving secondary stress were coded 2 (see Table S1). Unstressed syllables were coded 0. Despite the importance of stress information for spoken-language processing and its theoretical relevance to the TS framework, to our knowledge no previous study has directly examined the cortical encoding of intra-word prosodic prominence, either in typical development or in developmental dyslexia. Based on TS theory, we hypothesised here that adults with dyslexia would exhibit reduced cortical encoding of lexical stress patterns in the delta band.

Impaired cortical encoding of the envelope of continuous speech in the delta- and theta-bands has already been reported for adults with dyslexia (Keshavarzi et al., 2026b). It was studied here at the single word level. Two control metrics were computed, for word and syllable onsets. Word- and syllable-onset predictors mark discrete temporal boundaries but do not represent the continuous amplitude-modulation structure of speech or the relative prominence of syllables. Based on TS theory, we expected the cortical encoding of word and syllable onsets to be largely preserved. Cortical encoding was assessed independently for four complementary aspects of speech: the low-frequency speech envelope, word onset, syllable onset, and lexical stress pattern, using EEG and forward TRF modelling.

Participants with and without dyslexia listened to 101 naturally spoken English words and word-, syllable-, and stress-level predictors were derived using automated acoustic segmentation and forced phonetic alignment to obtain temporally precise annotations (Fig. 1a–d). To assess the behavioural relevance of any identified neural differences between groups, we examined whether cortical encoding measures were associated with individual differences in sight-word reading efficiency and phonemic decoding (nonword reading).

**Figure 1.**
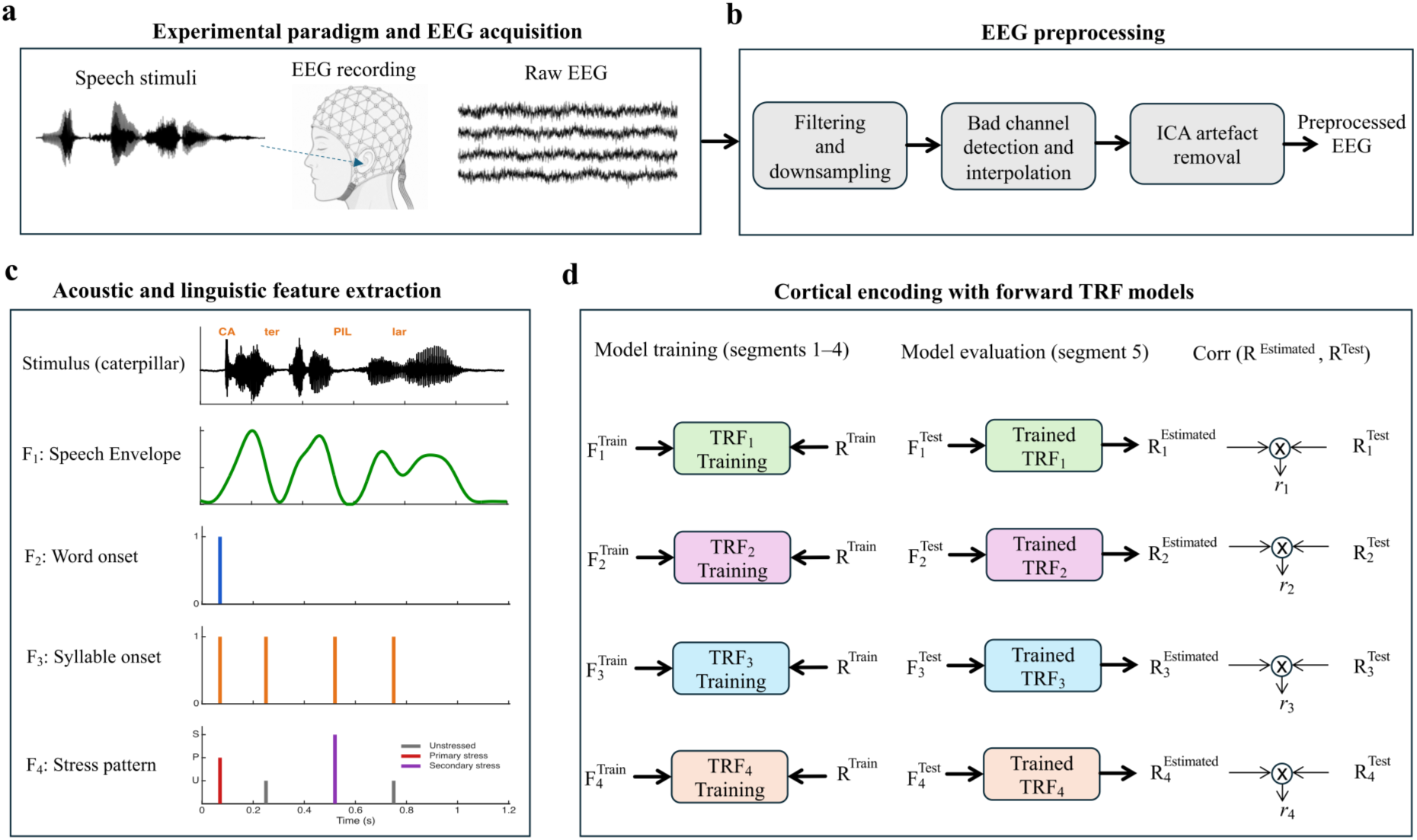
Experimental paradigm, EEG preprocessing, acoustic and linguistic feature extraction, and forward TRF modelling. (a) Participants listened to isolated spoken English words while EEG was recorded. (b) Continuous EEG data were filtered and downsampled, inspected for noisy channels that were corrected using bad-channel interpolation, and subsequently cleaned of artefacts using independent component analysis (ICA). (c) Four speech representations were extracted independently from the continuous speech recording: the low-frequency speech envelope, word onset, syllable onset, and lexical stress pattern. Word and syllable onsets were represented as binary impulse trains, whereas lexical stress was represented by three independent binary predictors corresponding to unstressed, primary-stressed, and secondary-stressed syllables. The word *caterpillar* is shown as an illustrative example. (d) Separate forward TRF models were trained using the first 80% (segments 1–4) of the stimulus recording and corresponding EEG data. Model performance was evaluated on the remaining 20% (segment 5) by predicting EEG responses from the held-out speech features. Encoding accuracy was quantified as the Pearson correlation between the predicted and recorded EEG signals and averaged across electrodes to obtain a single cortical encoding score for each participant and speech representation.

## Results

### Cortical encoding of the low-frequency speech envelope is reduced for those with dyslexia

As an initial validation, we first examined cortical encoding of the low-frequency speech envelope (0.5–8 Hz), which is known to be atypical regarding continuous speech processing from prior studies of those with dyslexia. This was done using forward TRF modelling in the delta (0.5–4 Hz) and theta (4–8 Hz) bands. As expected, adults with dyslexia exhibited significantly reduced cortical encoding of the speech envelope compared with typical readers for both frequency bands (Delta: *p* = 0.035; Theta: *p* = 0.012; Fig. 2a,b). Permutation analyses confirmed that cortical encoding exceeded chance levels for both groups for the delta-band (Control: *r* = 0.0596, *chance* = 0.0117; Dyslexia: *r* = 0.0338, *chance* = 0.0098; Supplementary Fig. S1a) and the group difference was significant (Fig. 2a). For the theta band, encoding exceeded chance for the control group (*r* = 0.0112, *chance* = 0.0070; Supplementary Fig. S1b) but not for the dyslexia group (*r* = −0.0033, *chance* = 0.0070) and the group difference was significant (Fig. 2b). Thus, adults with dyslexia showed reduced cortical representation of the low-frequency speech envelope for isolated words across both delta and theta timescales. This pattern is consistent with the predictions of TS theory. These results show that atypical low-frequency cortical speech encoding for those with dyslexia can occur in the absence of sentence- and discourse-level linguistic context.

**Figure 2.**
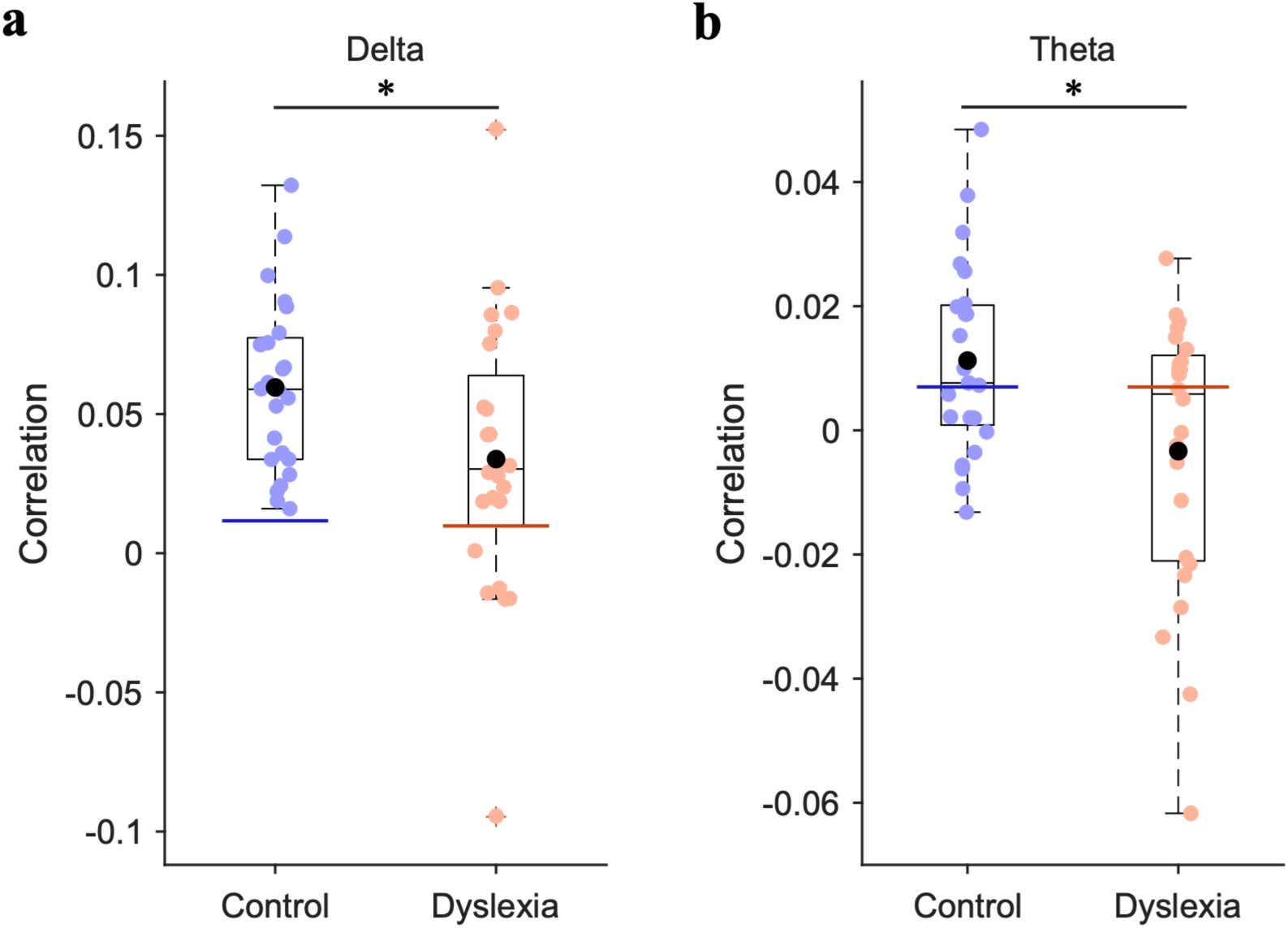
Cortical encoding of the low-frequency speech envelope in adults with and without dyslexia for the (a) delta and (b) theta bands. Box plots show the median, interquartile range, and whiskers extending to 1.5 × the interquartile range. Blue and orange points represent individual participants in the control and dyslexia groups, respectively. Black circles indicate group means. Horizontal blue and red lines indicate the 95th-percentile permutation-derived chance levels for the control and dyslexia groups, respectively. Asterisks indicate significant between-group differences (*p* < 0.05). Adults with dyslexia showed reduced cortical encoding of the speech envelope in both bands.

### Cortical encoding of lexical stress patterns is selectively impaired for those with dyslexia

We next examined cortical encoding of lexical stress patterns. To our knowledge, encoding of the relative prominence of syllables within a multisyllabic spoken word has not been studied previously using neural measures. A significant group difference was observed only for the delta band, adults with dyslexia showing lower lexical-stress encoding than controls (*p* = 0.031; Fig. 3a). Permutation analyses demonstrated that delta-band lexical-stress encoding exceeded the chance level for both groups (Control: *r* = 0.0324, *chance* = 0.0092; Dyslexia: *r* = 0.0151, *chance* = 0.0085; Supplementary Fig. S2a). This indicates measurable cortical encoding of lexical stress patterns for both groups, but with significantly weaker encoding for adults with dyslexia. In contrast, theta-band lexical-stress encoding did not differ significantly between groups (*p* = 0.542; Fig. 3b) and did not exceed the chance level for either group (Supplementary Fig. S2b). Thus, lexical-stress encoding appears to be confined to the delta band, those with dyslexia showing atypical cortical representation of prosodic prominence at a slow temporal timescale. Importantly, this effect was evident without the prosodic and contextual structure that is provided by continuous speech.

**Figure 3.**
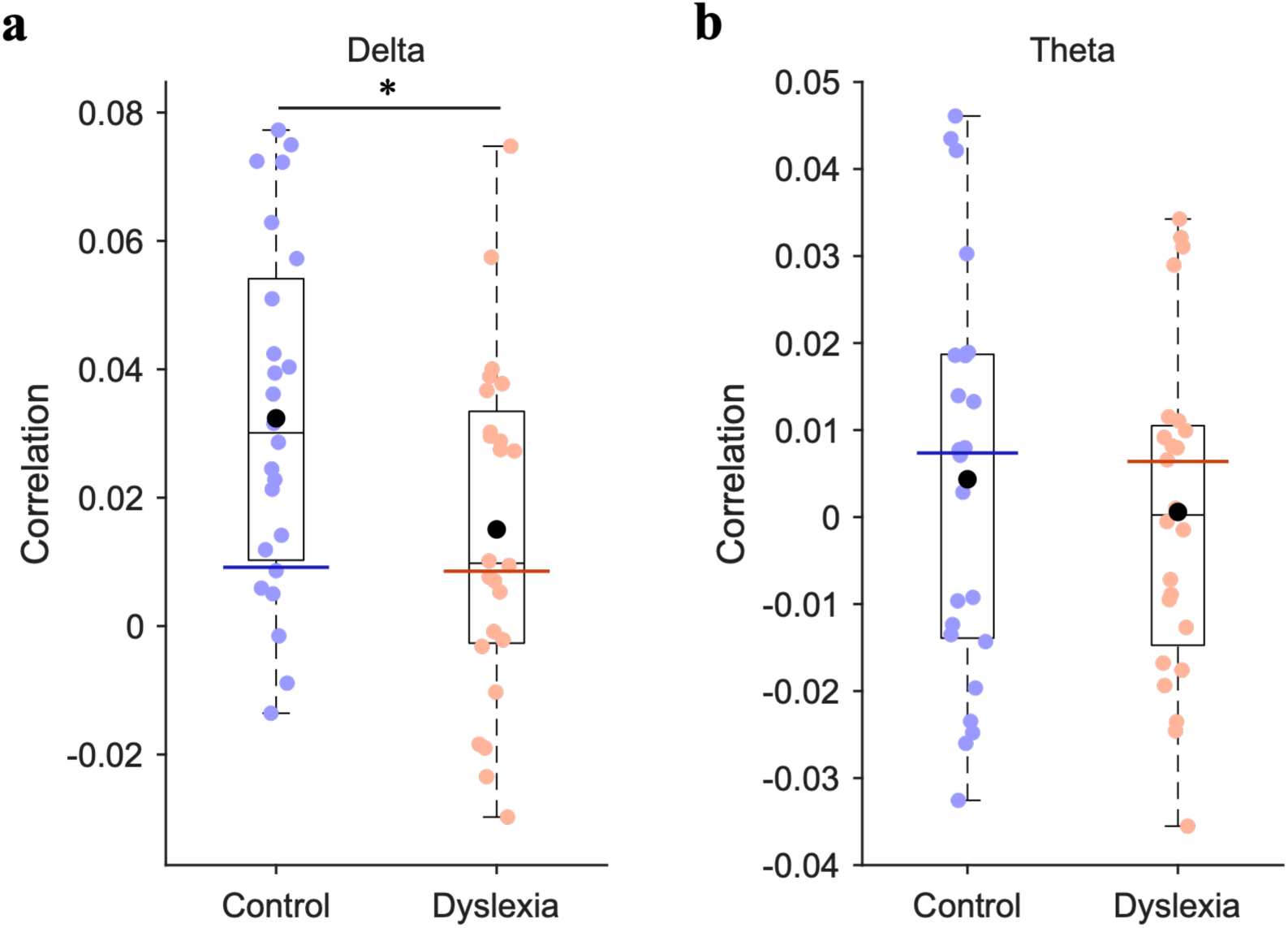
Cortical encoding of lexical stress patterns for adults with and without dyslexia for the (a) delta and (b) theta bands. Box plots show the median, interquartile range, and whiskers extending to 1.5 × the interquartile range. Blue and orange points represent individual participants in the control and dyslexia groups, respectively. Black circles indicate group means. Horizontal blue and red lines indicate the 95th-percentile permutation-derived chance levels for the control and dyslexia groups, respectively. Asterisks indicate significant between-group differences (*p* < 0.05). Adults with dyslexia showed reduced delta-band cortical encoding of lexical stress patterns, whereas theta-band encoding was not significant for either group and did not differ between groups.

### Cortical encoding of word onset is preserved for those with dyslexia

We next examined cortical encoding of the temporal onset of each spoken word. No significant group difference was observed for either the delta (*p* = 0.108; Fig. 4a) or theta (*p* = 0.937; Fig. 4b) band. Permutation analyses demonstrated that delta-band word-onset encoding substantially exceeded chance levels for both groups (Control: *r* = 0.0444, *chance* = 0.0089; Dyslexia: *r* = 0.0275, *chance* = 0.0086; Supplementary Fig. S3a), indicating robust cortical representation of word-onset timing at this slower temporal scale for adults with and without dyslexia. In contrast, theta-band word-onset encoding did not exceed chance levels for either group (Supplementary Fig. S3b), indicating that reliable encoding of word-onset timing was confined to the delta band under the present experimental conditions. The absence of a significant group difference in the delta-band contrasts with the reduced delta-band encoding of both the speech envelope and lexical stress patterns for those with dyslexia. This may indicate that dyslexia-related differences at this temporal scale do not extend to all speech representations.

**Figure 4.**
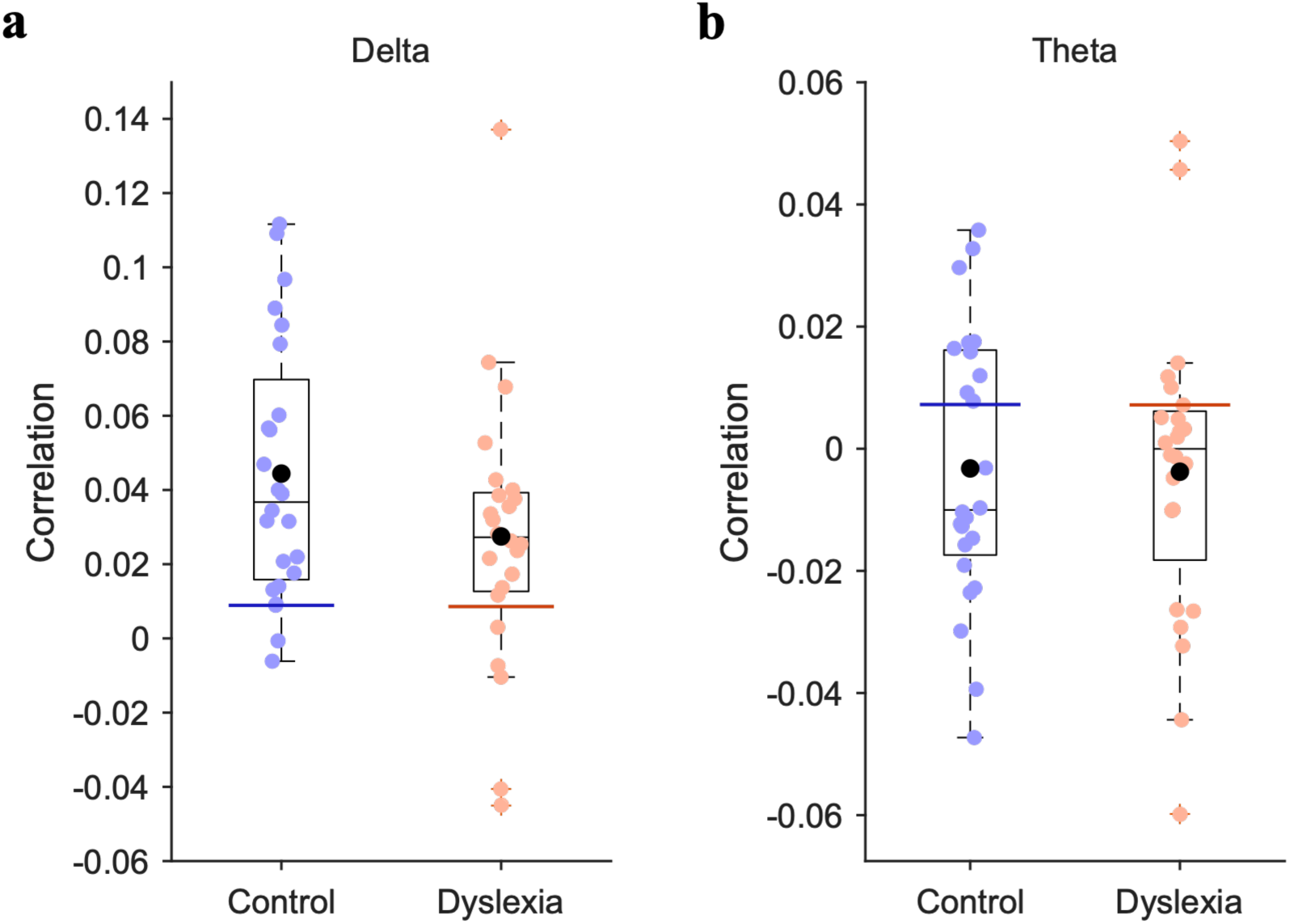
Cortical encoding of word onset for adults with and without dyslexia for the (a) delta and (b) theta bands. Box plots show the median, interquartile range, and whiskers extending to 1.5 × the interquartile range. Blue and orange points represent individual participants in the control and dyslexia groups, respectively. Black circles indicate group means. Horizontal blue and red lines indicate the 95th-percentile permutation-derived chance levels for the control and dyslexia groups, respectively. Cortical encoding of word onset did not differ between adults with dyslexia and control participants for either band.

### Cortical encoding of syllable onsets is preserved for those with dyslexia

Finally, we examined cortical encoding of syllable onsets. Similar to the word-onset analysis, there were no significant differences between groups for the delta (*p* = 0.825; Fig. 5a) and theta (*p* = 0.841; Fig. 5b) frequency bands. Both groups showed significant delta-band encoding relative to chance (Control: *r* = 0.0281, chance = 0.0082; Dyslexia: *r* = 0.0265, chance = 0.0073; Supplementary Fig. S4a). However, theta-band encoding was close to zero and did not exceed chance for either group (Supplementary Fig. S4b). Thus, cortical encoding of syllable-onset timing appears to be preserved in the delta band for those with dyslexia. This may indicate preserved cortical encoding of discrete temporal boundaries at the syllable level.

**Figure 5.**
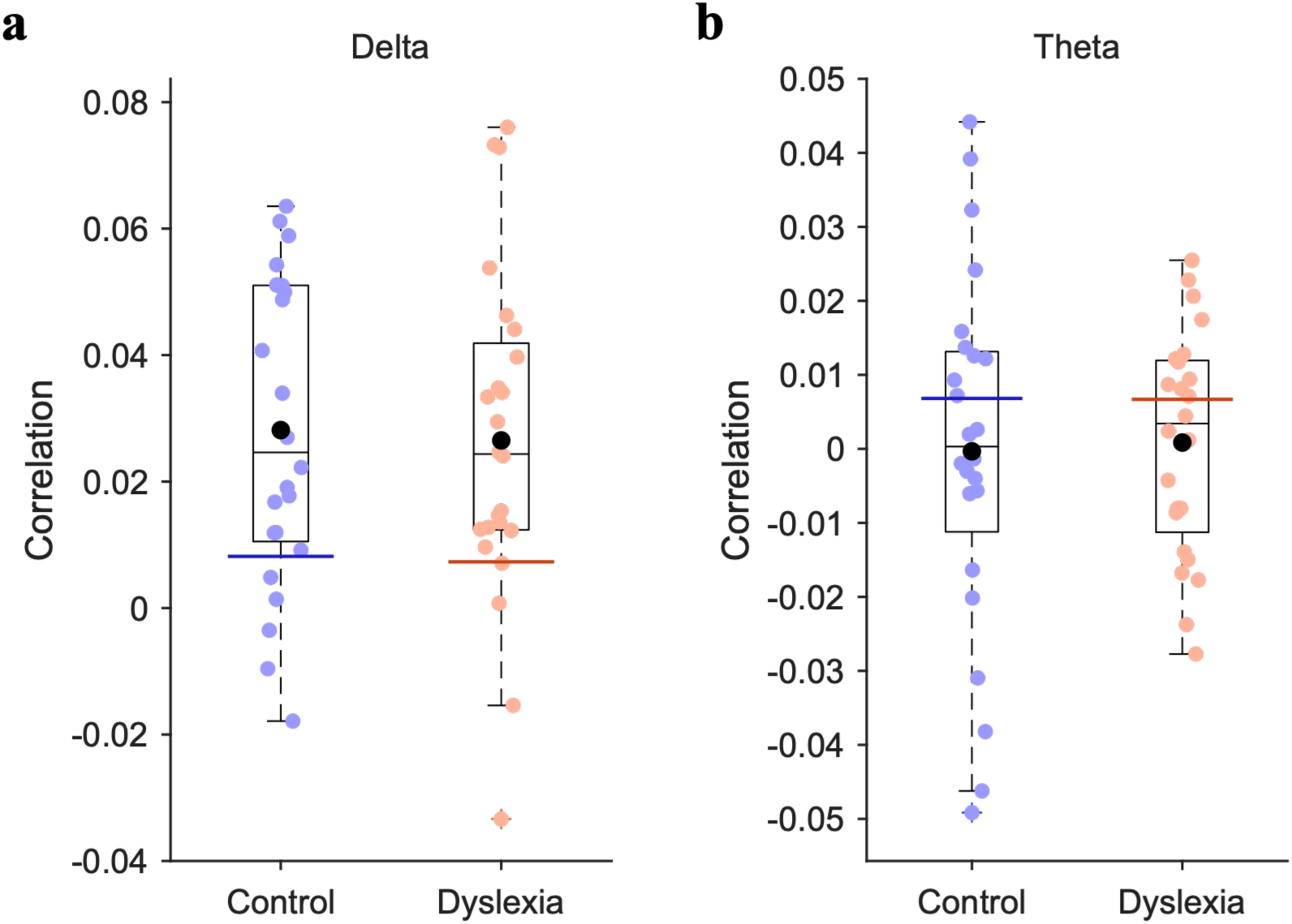
Cortical encoding of syllable onset for adults with and without dyslexia for the (a) delta and (b) theta bands. Box plots show the median, interquartile range, and whiskers extending to 1.5 × the interquartile range. Blue and orange points represent individual participants in the control and dyslexia groups, respectively. Black circles indicate group means. Horizontal blue and red lines indicate the 95th-percentile permutation-derived chance levels for the control and dyslexia groups, respectively. Cortical encoding of syllable onset did not differ between adults with dyslexia and control participants in either band and encoding was not significant for either group for the theta band.

### Cortical speech encoding is associated with phonemic decoding ability

To examine whether the cortical encoding measures showing significant group differences were associated with individual differences in reading ability (Fig. 6a–f), we assessed brain-behavioural correlations. Across all participants, stronger delta-band encoding of the low-frequency speech envelope was significantly associated with better phonemic decoding efficiency (TOWRE-PDE, a nonword reading measure; Spearman’s *ρ* = 0.41, false-discovery rate (FDR)-adjusted *p* = 0.022; Fig. 6b). However, the association between delta-band encoding and sight-word reading efficiency (TOWRE-SWE) was not significant (*ρ* = 0.26, FDR-adjusted *p* = 0.095; Fig. 6a). Theta-band speech-envelope encoding was not significantly associated with either sight-word reading efficiency (*ρ* = 0.13, FDR-adjusted *p* = 0.367; Fig. 6c) or phonemic decoding efficiency (*ρ* = 0.27, FDR-adjusted *p* = 0.095; Fig. 6d). Stronger delta-band encoding of lexical stress patterns was significantly associated with better phonemic decoding efficiency (*ρ* = 0.37, FDR-adjusted *p* = 0.032; Fig. 6f), but not with sight-word reading efficiency (*ρ* = 0.26, FDR-adjusted *p* = 0.095; Fig. 6e). Thus, stronger delta-band cortical encoding of both lexical stress patterns and the speech envelope was associated with better nonword reading, a task that is thought to draw heavily on phonological processes. This pattern is consistent with the TS framework.

**Figure 6.**
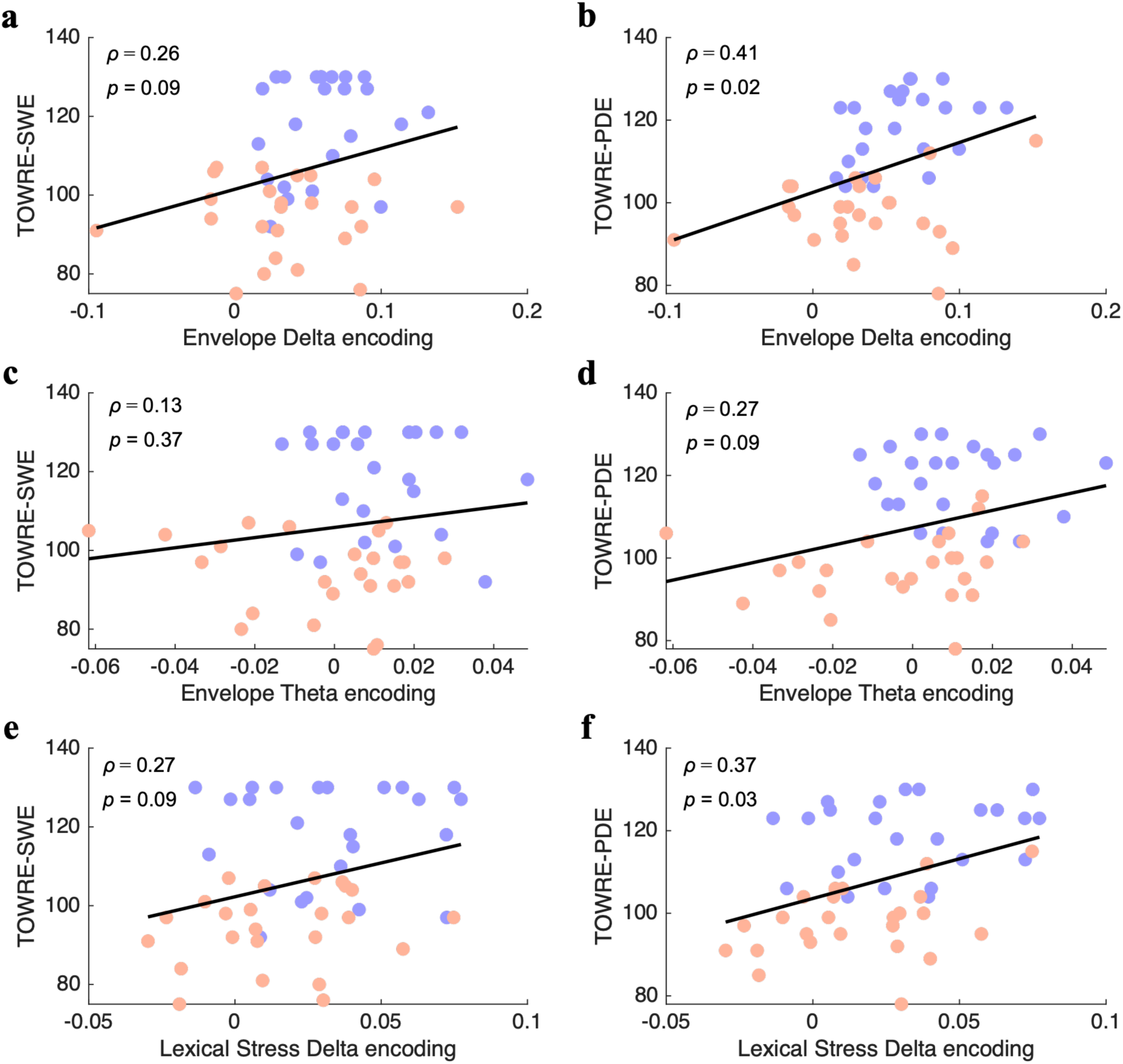
Associations between cortical speech encoding and reading ability scores: (a) delta-band speech-envelope encoding and TOWRE-SWE, (b) delta-band speech-envelope encoding and TOWRE-PDE, (c) theta-band speech-envelope encoding and TOWRE-SWE, (d) theta-band speech-envelope encoding and TOWRE-PDE, (e) delta-band lexical-stress encoding and TOWRE-SWE, and (f) delta-band lexical-stress encoding and TOWRE-PDE. Scatterplots show associations between cortical encoding accuracy and standardised measures of sight-word reading efficiency (TOWRE-SWE) and phonemic decoding efficiency (TOWRE-PDE) across participants. Blue and orange points represent individual control and dyslexic participants, respectively. Black lines show least-squares linear fits for visualisation; statistical associations were assessed using Spearman’s correlations. Spearman’s *ρ* and FDR-adjusted *p* values are reported for each association. Stronger delta-band encoding of both the speech envelope (b) and lexical stress patterns (f) was significantly associated with better phonemic decoding efficiency.

## Discussion

The present study investigated the question of whether atypical cortical encoding of prosodic prominence is evident in adults with developmental dyslexia during isolated multisyllabic word perception. Cortical encoding was also examined in the delta and theta bands for the low-frequency speech envelope, word onset, and syllable onset. By using isolated spoken words, we substantially reduced the higher-level linguistic and cognitive demands associated with the processing of continuous natural speech, including syntactic integration, contextual prediction, discourse-level processing and working memory. The results revealed a selective pattern of impairment. While group differences were evident for representations of prosodic prominence and continuous acoustic modulation (the speech envelope), cortical encoding of word onset timing and syllable onset timing did not differ significantly between groups. This dissociation between relatively preserved word and syllable onset encoding and reduced speech-envelope and lexical-stress encoding argues against a general reduction in cortical sensitivity to the timing of speech events for those with dyslexia. Instead, the findings indicate more selective group differences in the cortical representation of slow acoustic modulations and prosodic structure, consistent with TS theory.

Reduced cortical encoding of the speech envelope for single multisyllabic words was found for both the delta and theta bands. This is of interest, as these timescales have been proposed to support complementary aspects of speech processing. Slower delta-timescale activity is particularly relevant to the coding of prosodic and phrasal structure and theta-timescale activity is closely associated with the coding of syllabic-rate information (Giraud & Poeppel, 2012). Reduced envelope encoding across both bands is consistent with atypical cortical sensitivity to temporal speech structure across multiple low-frequency timescales. Note that theta-band envelope encoding exceeded the permutation-derived chance threshold for control adults but not for adults with dyslexia. Failure to exceed an empirical chance threshold does not provide evidence for the absence of a neural representation. However, when considered alongside the significant between-group difference, it appears to provide converging evidence for particularly weak theta-band representation of the word-level speech envelope for adults with dyslexia.

The significant reduction in delta-band cortical encoding of intra-word prosodic prominence for those with dyslexia provides direct neurophysiological support for the speech modelling studies based on TS theory (Leong et al., 2014; Leong & Goswami, 2015; Leong et al., 2017). These studies indicated that phase relations dependent on delta-rate amplitude modulations in the speech envelope governed prosodic prominence, as peak-to-peak matching of delta- and theta-rate amplitude modulations was required for the perception of stressed syllables. The current data show that cortical representation of within-word prosodic prominence depends on delta-rate neural encoding. In the neural oscillatory hierarchy, delta phase governs theta phase (Gross et al., 2013). These adult impairments are interesting, as children with both dyslexia (Keshavarzi et al., 2024) and DLD (Parvez et al., 2024) find it difficult to copy within-word prosodic prominence and they also have delta-band cortical speech tracking impairments (Keshavarzi et al., 2022a; 2026b). It is difficult to produce aspects of phonology that you cannot discriminate very well.

The brain–behaviour associations reported for nonword reading support the functional relevance of these neural differences. A deficit in nonword reading is the primary hallmark of developmental dyslexia across languages. Across all participants, stronger delta-band lexical-stress encoding was associated with better phonemic decoding efficiency, and stronger delta-band speech-envelope encoding showed a similar relationship. The association of phonemic decoding with both delta-band envelope and lexical-stress encoding is consistent with TS theory, which focuses on the development of the phonological system from birth (Goswami, 2011, 2022b; Di Liberto et al., 2023). According to TS theory, atypical neural representation of slow speech modulations and prosodic structure contributes to the development of less precise phonological representations at every linguistic level (lexical stress, syllable, rhyme and phoneme) from infancy onwards, adversely affecting the acquisition of reading for all languages.

Several limitations of this study should be considered. First, although isolated words substantially reduce sentence- and contextual processing, they do not provide a purely bottom-up measure of auditory processing. Individual words engage lexical access, phonological knowledge and potentially predictive processes. The present findings should therefore be interpreted as demonstrating atypical cortical encoding in the absence of continuous linguistic context, rather than as isolating exclusively bottom-up neural mechanisms. Second, the present sample comprised university students with dyslexia and may therefore represent relatively high-functioning adults who have developed substantial compensatory strategies. Examining lexical-stress encoding longitudinally for children who are learning to read will be important for determining whether the neural differences associated with dyslexia precede reading difficulty or develop alongside it. Third, the observed brain–behaviour associations were calculated across the full sample of participants and may reflect between-group differences in both cortical encoding and reading performance. The associations should not be interpreted as demonstrating a group-independent or causal relationship. Future studies with larger samples could examine these relationships within groups and across development.

In conclusion, the present study demonstrates that atypical low-frequency cortical speech encoding for adults with developmental dyslexia is evident during the perception of isolated multisyllabic words, with reduced delta-band encoding related to the representation of prosodic prominence and reduced speech-envelope encoding for both the delta and theta bands. In contrast, cortical encoding of word- and syllable-onset timing was preserved. Moreover, stronger delta-band encoding of both the speech envelope and lexical stress patterns was associated with better nonword reading, a core difficulty for those with developmental dyslexia. This selective reduction in the cortical representation of slow acoustic and prosodic speech information, alongside relatively preserved encoding of discrete speech-onset timing, provides further neurophysiological support for TS theory (Goswami, 2011).

## Materials and Methods

### Participants

Forty-eight adult native English speakers participated in the study, including 24 control participants (mean age = 21.9 years, SD = 2.6; range = 18.7–29.2 years) and 24 adults with dyslexia (mean age = 21.4 years, SD = 2.5; range = 18.3–29.1 years). All participants were university students. Participants with dyslexia were recruited through the Accessibility & Disability Resource Centre at the University of Cambridge and had a formal diagnosis of dyslexia. Inclusion and exclusion criteria were specified before data analysis. Dyslexic participants were required to have no additional reported neurodevelopmental or learning conditions including autism spectrum disorder, dyspraxia, attention deficit hyperactivity disorder, or developmental language disorder. The absence of the additional conditions was established by self-report. Dyslexic participants were also required to score at least one standard deviation below the control-group mean on at least one subtest of the Test of Word Reading Efficiency–Second Edition (TOWRE-2; Torgesen et al., 2012). The control group was matched as closely as possible to the dyslexia group for age and nonverbal reasoning ability, assessed using the Matrix Reasoning subtest (Wechsler, 2008; Table 1). There were no significant group differences in age or Matrix Reasoning performance. As expected, the group with dyslexia performed significantly more poorly on the Sight Word Efficiency and Phonemic Decoding Efficiency subtests of the TOWRE-2. Mathematical ability, as assessed using the Test of Written Arithmetic (TAN; Wilkinson & Robertson, 2006), did not differ significantly between groups. Pure-tone hearing thresholds were measured bilaterally at 250, 500, 1000, 2000, 4000, and 8000 Hz. All participants had thresholds of 20 dB HL or better at every test frequency. The same participant cohort was included in Keshavarzi (2025) and Keshavarzi et al. (2026b, 2026c). All participants provided written informed consent in accordance with the Declaration of Helsinki. The study was reviewed by the Psychology Research Ethics Committee at the University of Cambridge and received a favourable opinion.

**Table 1.** Participant characteristics and behavioural performance for the control and dyslexia groups. Values are means with standard deviations in parentheses. Measures included age and scores for TOWRE-SWE, TOWRE-PDE, Matrix Reasoning, and TAN. Group differences were evaluated using two-tailed Wilcoxon rank-sum tests.

|  | Control | Dyslexia | Test outcome |
| --- | --- | --- | --- |
| **Age (year)** | 21.9 (2.6) | 21.4 (2.5) | $p = 0.55$ |
| **TOWRE-SWE** | 118 (12.9) | 94 (9.7) | $p = 1.1 \times 10^{-6}$ |
| **TOWRE-PDE** | 119 (8.9) | 98 (8.3) | $p = 5.8 \times 10^{-8}$ |
| **Matrix Reasoning** | 15 (1.6) | 15 (1.8) | $p = 0.53$ |
| **TAN** | 76 (10) | 72 (11.6) | $p = 0.22$ |

### Experimental setup and stimuli

Participants were seated in an electrically shielded, sound-attenuated room to minimise environmental noise and electromagnetic interference. Auditory stimuli were presented binaurally through ER-2 insert earphones (Etymotic Research) at a sampling rate of 44.1 kHz. EEG was recorded concurrently at 1000 Hz using a 128-channel HydroCel Geodesic Sensor Net (Electrical Geodesics). The stimuli comprised 101 naturally spoken English words (95 unique words, of which 6 were repeated) produced by a female native speaker of British English, which varied in syllable number and lexical stress pattern and were recorded as a single continuous audio file, with silent inter-word gaps ranging from approximately 0.9 to 3.1 s (median 1.9 s). As part of the broader PhD paradigm, six words were repeated within the stimulus sequence to permit a secondary assessment of within-session response consistency. These repetitions were not analysed separately in the present study. Participants were instructed to remain still and fixate on a red cross displayed at the centre of a monitor to minimise eye-movement artefacts while listening to the stimuli. EEG was recorded continuously throughout stimulus presentation.

### EEG preprocessing

EEG data were initially referenced to Cz and band-pass filtered between 0.2 and 42 Hz using a zero-phase finite impulse response filter. The data were then downsampled to 200 Hz. Noisy channels were identified by visual inspection of their temporal and spectral characteristics and were interpolated using MNE-Python (Gramfort et al., 2013). Independent component analysis was performed using the Infomax algorithm. Components reflecting eye blinks, eye movements, cardiac activity, or other non-neural artefacts were identified from their scalp topographies, power spectra, and time-domain waveforms and were removed before further analysis. The preprocessed EEG was filtered separately into the delta (0.5–4 Hz) and theta (4–8 Hz) bands using third-order Butterworth high-pass and low-pass filters applied in both temporal directions to achieve zero-phase filtering. The EEG was subsequently downsampled from 200 to 100 Hz.

### Acoustic and linguistic feature extraction

Four acoustic and linguistic features were extracted: the low-frequency speech envelope, word onset, syllable onset, and lexical stress pattern. Feature extraction was performed using an automated processing pipeline developed for the present study, combining acoustic segmentation with forced phonetic alignment to obtain temporally precise annotations of each speech feature. The low-frequency speech envelope was obtained as the magnitude of the analytic signal computed using the Hilbert transform of the broadband speech waveform, low-pass filtered at 50 Hz, and resampled to 200 Hz.

Word onset and offset times were identified automatically using an amplitude-based segmentation procedure. The amplitude envelope of the recording was computed as the magnitude of the analytic signal obtained using the Hilbert transform of the broadband waveform. Candidate speech-active intervals were defined as continuous regions in which the envelope exceeded a threshold 30-dB below the maximum amplitude of the recording. Brief intervals shorter than 40 ms were excluded, and adjacent speech-active intervals separated by less than 700 ms were merged, yielding 101 word tokens. The 700-ms merge criterion was chosen to bridge brief within-word amplitude dips (e.g., stop-consonant closures) without merging separate words, based on inspection of the inter-word pause distribution in this recording. The onset and offset times of each token were subsequently refined using a local amplitude threshold corresponding to 3% of the token peak amplitude, within 150-ms search windows preceding the initially identified onset and following the initially identified offset, respectively.

Syllable onset times and lexical stress patterns were obtained using forced phonetic alignment. Each word token was aligned at the phoneme level with the English (US) ARPA acoustic model and pronunciation dictionary, using the Montreal Forced Aligner (MFA; McAuliffe et al., 2017). Syllable boundaries were determined from the resulting phoneme alignments using a maximal-onset syllabification algorithm, whereby intervocalic consonants were assigned to the onset of the following syllable. The lexical stress category assigned to each syllable (primary, secondary, or unstressed) was obtained from the stress digit associated with its vowel nucleus in the pronunciation dictionary.

For encoding modelling, the word-onset and syllable-onset features were each represented as univariate impulse trains, with impulses (value = 1) marking the onset of each event and zeros elsewhere. Lexical stress pattern was represented categorically as three simultaneous binary impulse trains corresponding to unstressed, primary-stress, and secondary-stress syllable onsets. This representation avoided imposing the assumption that neural responses vary linearly with stress level and allowed each stress category to contribute independently to the encoding model. All stimulus features were computed at a sampling rate of 200 Hz and time-aligned to the same trial structure as used for the speech envelope and EEG data. All features were subsequently downsampled from 200 to 100 Hz.

### Forward temporal response function (TRF) modelling

Cortical encoding was quantified using forward TRF modelling implemented in the multivariate TRF Toolbox (Crosse et al., 2016). Forward TRF models estimate the EEG response from one or more speech features over a range of temporal lags spanning 0–400 ms post-stimulus, thereby characterising how individual speech representations are encoded by the cortex over this time window. Separate encoding models were constructed for each participant, frequency band (delta or theta), and speech feature.

Four feature sets were modelled independently: the low-frequency speech envelope, word onset, syllable onset, and lexical stress pattern. The envelope model contained a single continuous predictor. The word-onset and syllable-onset models each contained a single binary impulse-train predictor. The lexical-stress model contained three simultaneous binary predictors representing unstressed, primary-stressed, and secondary-stressed syllable onsets. Each feature set was analysed separately, and no combined-feature models were fitted.

Before TRF modelling, EEG and stimulus features were temporally aligned and resampled to 100 Hz. The speech-envelope predictor was band-pass filtered between 0.5 and 8 Hz before resampling. In contrast, the onset-based predictors were not filtered because they consisted of discrete impulse events. During downsampling, these predictors were resampled using maximum-value pooling to preserve the precise timing of onset events. EEG amplitudes were normalised by the standard deviation calculated across all channels, samples, and trials for each participant, and each stimulus predictor was normalised independently within each trial.

The recording of the words was divided into five segments. The first four segments were used for model training, whereas the fifth segment (20% of the recording) served as an independent test set. The optimal ridge-regularisation parameter (λ) was selected using leave-one-segment-out cross-validation performed on the four training segments. Eight candidate λ values, ranging from 10^−1^ to 10^6^, were evaluated, and the value yielding the highest mean prediction accuracy across the four cross-validation folds and all EEG channels was selected. A final TRF model was then trained using the complete training data and evaluated on the independent fifth segment. Model performance was quantified as the Pearson correlation between the predicted and recorded EEG signals for each of the 128 electrodes. Correlation coefficients were then averaged across all electrodes, as implemented in the mTRF Toolbox, to obtain a single measure of cortical encoding for each participant. These participant-level encoding scores were used for all subsequent statistical analyses. Separate encoding scores were obtained for each speech feature and frequency band.

### Chance-level permutation analysis

Chance-level cortical encoding accuracy was estimated separately for each participant, frequency band, and speech feature using a permutation-based null model. For each of 500 permutations, the stimulus representation was circularly shifted independently within each stimulus segment while the corresponding EEG data remained unchanged. This procedure disrupted the temporal correspondence between the stimulus representation and neural response while preserving the temporal structure and autocorrelation of the stimulus predictor. The shift magnitude was randomly sampled for each segment subject to a predefined minimum offset. For the continuous speech-envelope predictor, a minimum shift of 2 s was used, corresponding to one full cycle at the lowest frequency retained in the envelope representation (0.5 Hz). For the impulse-based word-onset, syllable-onset, and lexical-stress predictors, a minimum shift of 0.5 s was used.

For each permutation, the temporally shifted stimulus representations from segments 1–4 were used to train a null forward TRF model, and encoding accuracy was evaluated on the shifted stimulus representation from the same held-out fifth segment as used in the empirical analysis. The ridge-regularisation parameter (λ) was not re-estimated for individual permutations. Instead, for each participant, frequency band, and feature set, λ was selected once by cross-validation using the original temporally aligned training data and subsequently held fixed across all 500 permutations. This ensured that the empirical and null models had the same regularisation strength while avoiding re-optimisation of model complexity for temporally misaligned stimulus representations.

For each permutation, prediction accuracy was quantified as the Pearson correlation between predicted and observed EEG responses for each of the 128 channels and then averaged across channels, as implemented in the mTRF Toolbox, to yield a single participant-level null encoding score. Group-level null distributions were constructed separately for the control group, group with dyslexia, and full sample by averaging participant-level null scores across the relevant participants at each permutation. This yielded 500 group-level null values for each speech feature and frequency band. The 95th percentile of each null distribution was defined as the empirical chance-level threshold and was compared with the corresponding observed group-mean encoding accuracy.

### Brain–behaviour correlation analysis

To examine the behavioural relevance of cortical speech encoding, brain–behaviour associations were assessed for neural measures that showed significant differences between the control group and group with dyslexia in the primary TRF analyses. These comprised delta- and theta-band encoding of the low-frequency speech envelope and delta-band encoding of lexical stress patterns. For each neural measure, the correlation was determined between encoding accuracy and standardised measures of sight-word reading efficiency (TOWRE-SWE) and phonemic decoding efficiency (TOWRE-PDE) from the Test of Word Reading Efficiency–Second Edition (TOWRE-2; Torgesen et al., 2012). Associations were assessed across all participants using two-tailed Spearman rank correlations. Six brain–behaviour correlations were evaluated in total, and p values were corrected for multiple comparisons across these six tests using the Benjamini–Hochberg false discovery rate (FDR) procedure, with statistical significance defined as FDR-adjusted *p* < 0.05.

## Author contributions

M.K. conceptualisation, methodology, data collection, data analyses, visualisation, writing-original draft; B.C.J.M. conceptualisation, methodology, writing-review & editing; U.G. funding acquisition, project administration, supervision, conceptualisation, methodology, writing-original draft.

## Acknowledgements

The authors would like to thank all the participants involved in the study. This research was funded by a donation to UG from the Yidan Prize Foundation. The sponsor played no role in the study design, data interpretation nor writing of the report.

## Supplementary Information

**Table S1.**
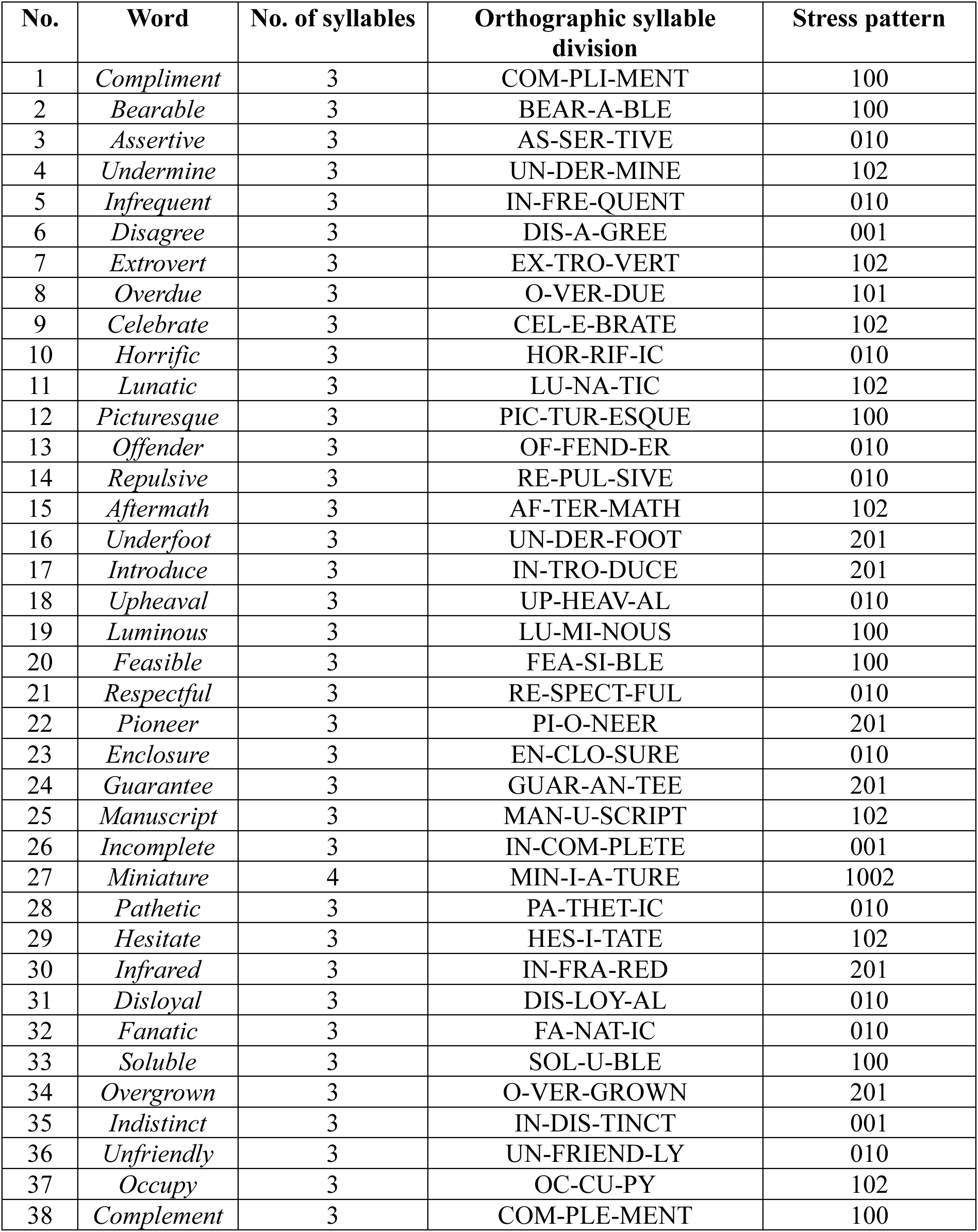
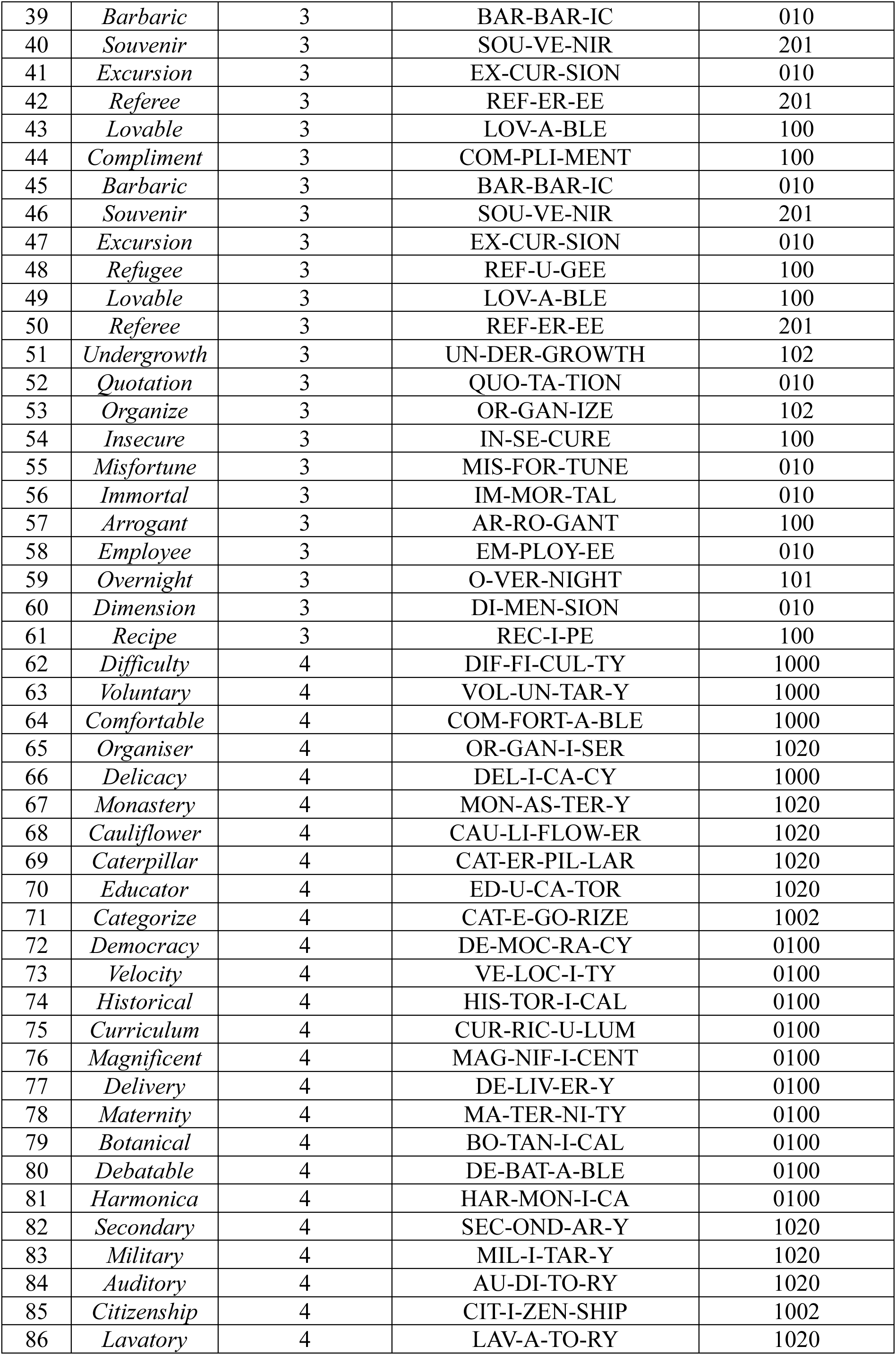
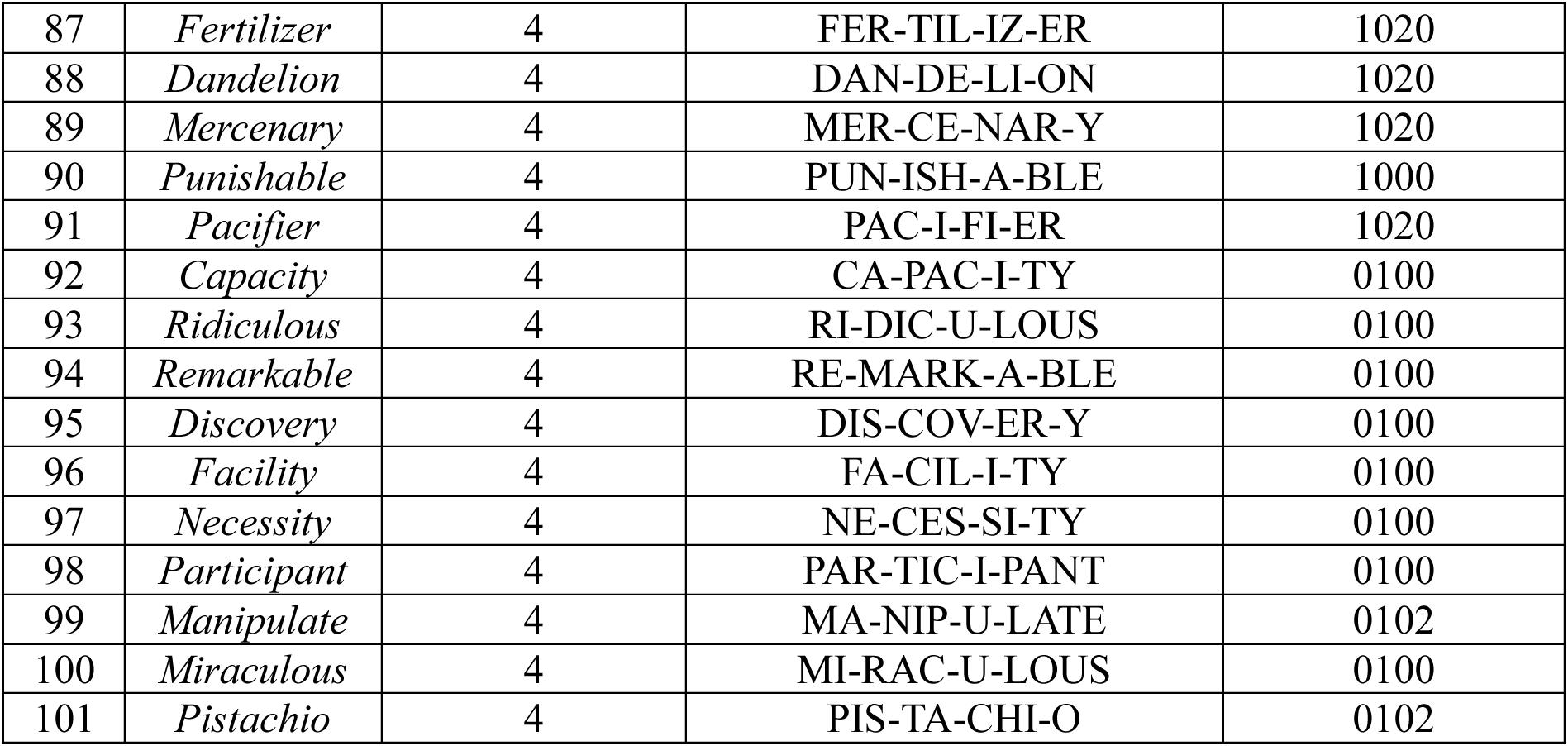
Full stimulus word list with syllable and lexical-stress coding. All 101 word tokens presented during the experiment are listed in presentation order (95 unique words, of which 6 were repeated), with syllable count, orthographic syllable division, and lexical-stress pattern (1 = primary stress, 2 = secondary stress, 0 = unstressed syllable), following the coding convention described in the main text.

**Figure S1.**
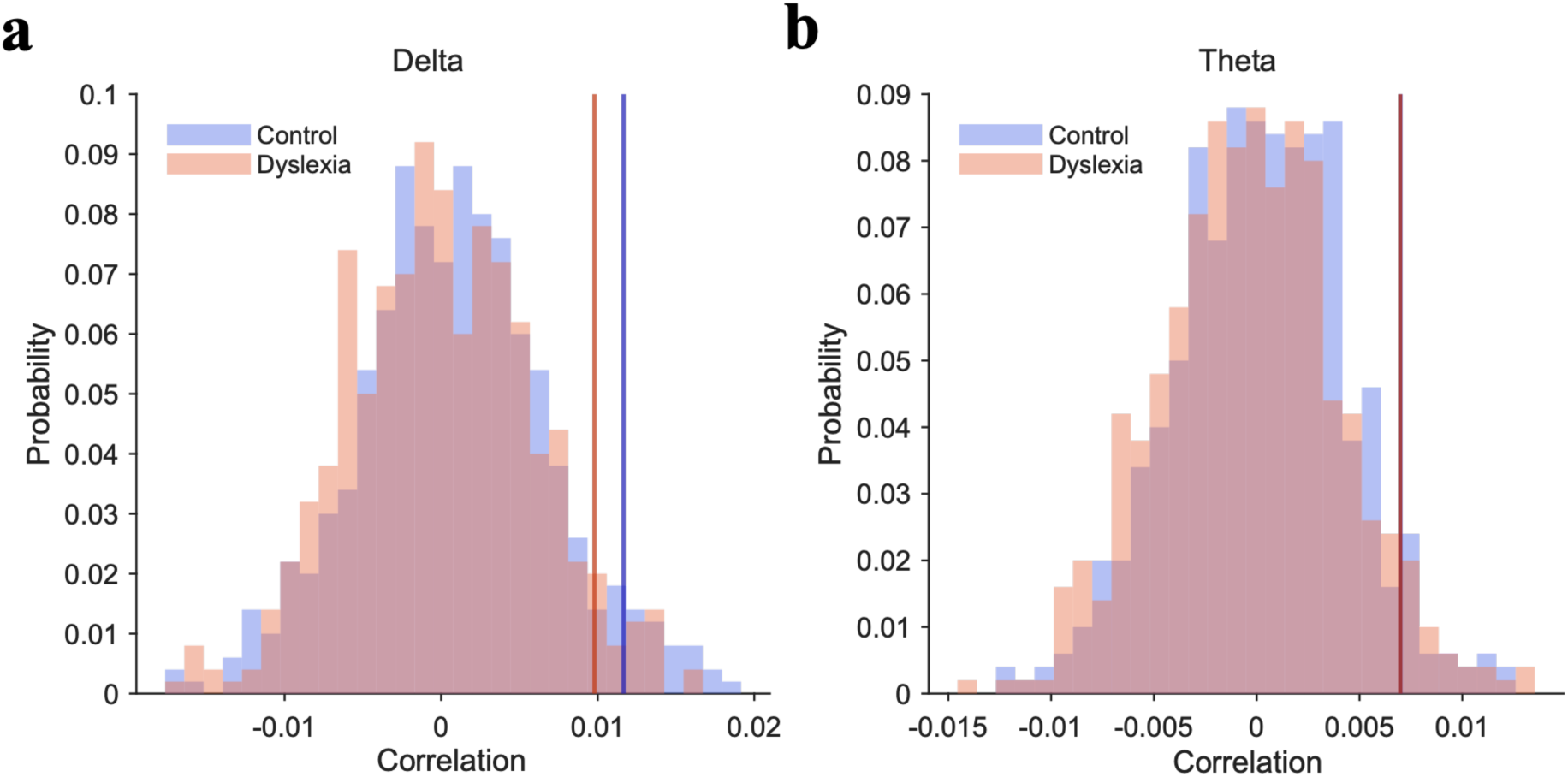
Chance-level distributions of correlations for speech-envelope encoding. Permutation-derived null distributions of TRF encoding accuracy for the speech-envelope predictor in the (a) delta and (b) theta bands. Blue distributions represent the control group and orange distributions represent the group with dyslexia. Blue and orange vertical lines indicate the corresponding group-specific 95th-percentile chance thresholds.

**Figure S2.**
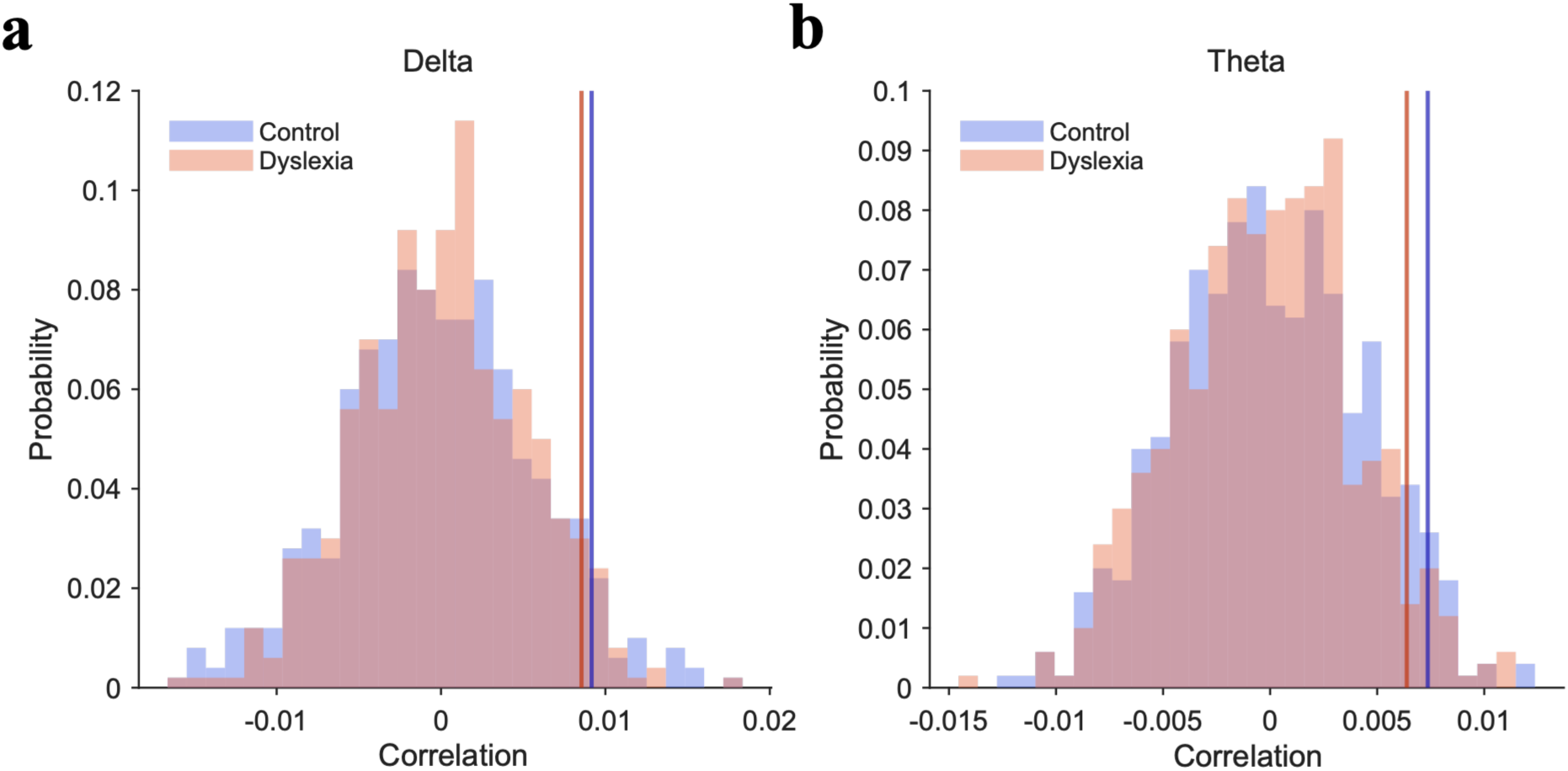
Chance-level distributions of correlations for lexical-stress encoding. Permutation-derived null distributions of TRF encoding accuracy for the lexical-stress predictor in the (a) delta and (b) theta bands. Blue distributions represent the control group and orange distributions represent the group with dyslexia. Blue and orange vertical lines indicate the corresponding group-specific 95th-percentile chance thresholds.

**Figure S3.**
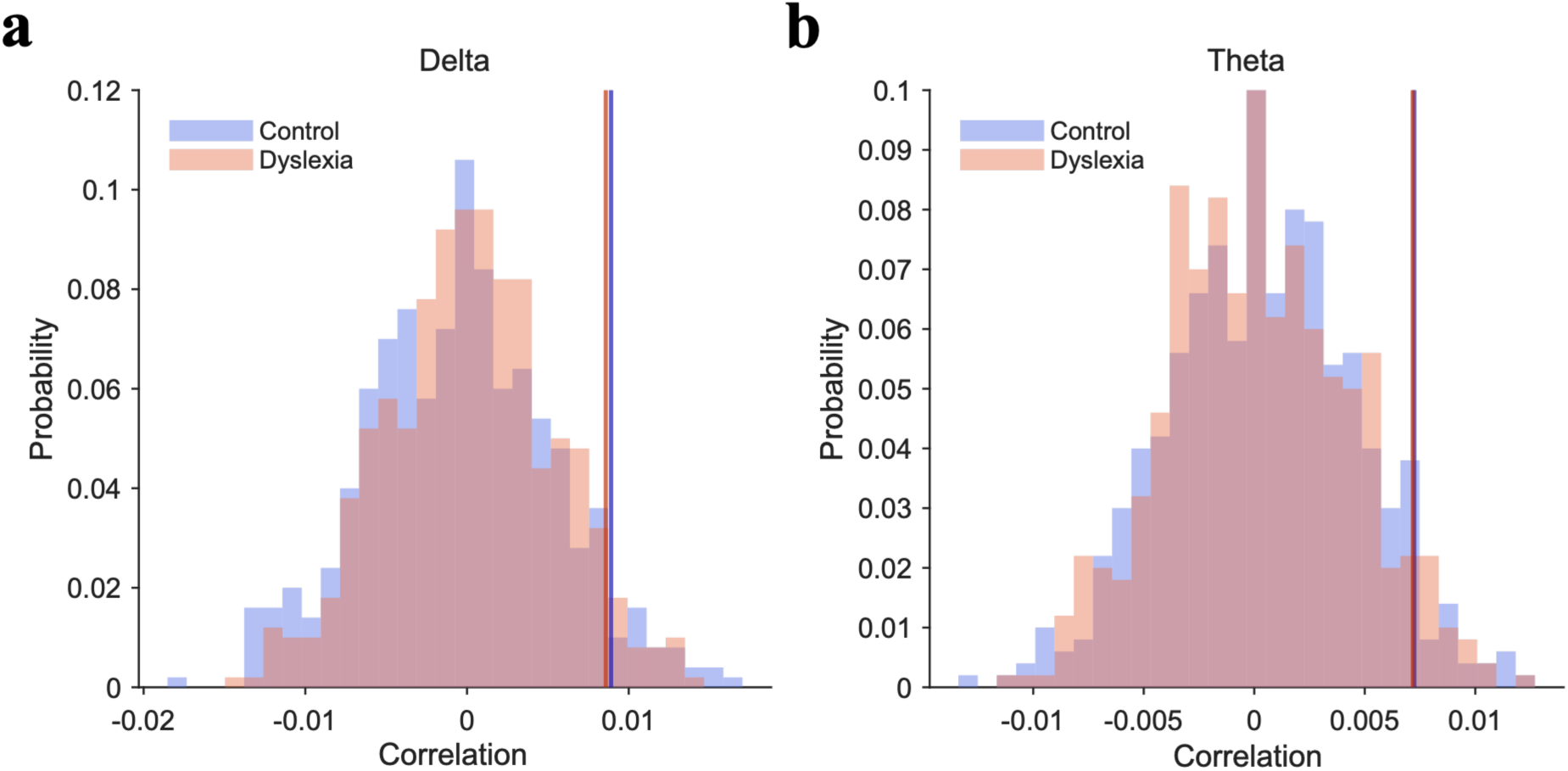
Chance-level distributions of correlations for word-onset encoding. Permutation-derived null distributions of TRF encoding accuracy for the word-onset predictor in the (a) delta and (b) theta bands. Blue distributions represent the control group and orange distributions represent the group with dyslexia. Blue and orange vertical lines indicate the corresponding group-specific 95th-percentile chance thresholds.

**Figure S4.**
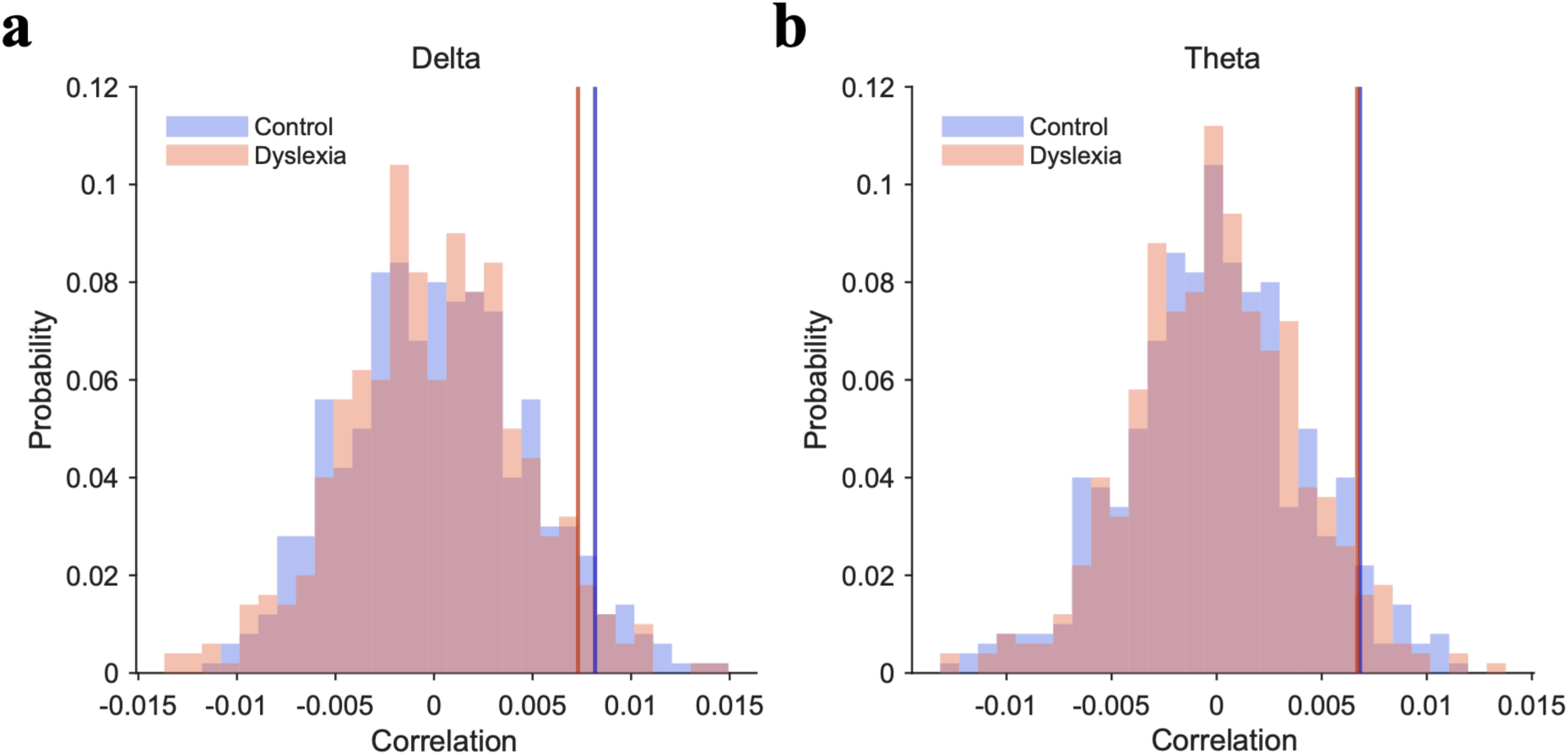
Chance-level distributions of correlations for syllable-onset encoding. Permutation-derived null distributions of TRF encoding accuracy for the syllable-onset predictor in the (a) delta and (b) theta bands. Blue distributions represent the control group and orange distributions represent the group with dyslexia. Blue and orange vertical lines indicate the corresponding group-specific 95th-percentile chance thresholds.

